# A Proposal of New Feature Selection Method Sensitive to Outliers and Correlation

**DOI:** 10.1101/2021.03.11.434934

**Authors:** Mert Demirarslan, Aslı Suner

**Affiliations:** Ege University, Faculty of Medicine, Department of Biostatistics and Medical Informatics, İzmir, Turkey

**Keywords:** Missing Value, Class Noise Class Imbalanced Outliers, Feature Selection, Ensemble Learning

## Abstract

In disease diagnosis classification, ensemble learning algorithms enable strong and successful models by training more than one learning function simultaneously. This study aimed to eliminate the irrelevant variable problem with the proposed new feature selection method and compare the ensemble learning algorithms’ classification performances after eliminating the problems such as missing observation, classroom noise, and class imbalance that may occur in the disease diagnosis data. According to the findings obtained; In the preprocessed data, it was seen that the classification performance of the algorithms was higher than the raw version of the data. When the algorithms’ classification performances for the new proposed advanced t-Score and the old t-Score method were compared, the feature selection made with the proposed method showed statistically higher performance in all data sets and all algorithms compared to the old t-Score method (p = 0.0001).

## 1. INTRODUCTION

Artificial intelligence applications are widely used in healthcare. In the field of health, it is seen that classification algorithms are frequently used, especially in diagnosing diseases, since rapid decisions are required on vital issues [1]. Algorithms used in disease diagnosis data sets should contain accurate and high-performance values. Therefore, it is of great importance that the data used are neat, clean, and suitable for the use of classification algorithms. However, many problems may arise in data sets, such as missing observation, class noise, class imbalance, outlier observation, correlation, and irrelevant variables [2]. In this case, it may affect the performance values of the algorithms negatively.

One of the most common problems in health data is the **missing value** problem. The existence of missing observations adversely affects the algorithms’ working system, and the missing observation situation is generally not taken into account in the literature. Some assignment methods such as mean, median, and regression assignment solve the missing observation problem [3]. Liu et al. (2017) assigned the smallest value to missing values in their study [4]. Min Wei et al. (2018) assigned the missing values in health data sets using k-nearest neighbor, multi-layer sensors, and support vector machine method [5].

Another common problem in health data is the **class imbalance** problem. Considering the general population and the prevalence of diseases; In the data, the number of patients is less than the number of healthy people. Class imbalance causes algorithms to learn biased (in terms of healthy individuals) and give wrong results. It is recommended that the classes are balanced so that the algorithms work properly, do not bias and learn incorrectly. Leo et al. (2019) tried to solve the class imbalance problem with the Adaptive Synthetic Sampling Approach-ADASYN in their study [6]. Alghamdi et al. (2017) investigated the effect of the SMOTE (Synthetic Minority Over-sampling Technique) method on the classification performance of ensemble learning methods in their study for the diagnosis of diabetes. As a result, they made their data balanced by solving the unbalanced class problem with the SMOTE method. In their study, Hyun Seo and Hyuk Kim (2018) optimized the SMOTE algorithm’s parameters to achieve better classification performance in unstable data [7].

**The class noise** problem is not seen much in health data sets, but it is another problem that causes problems in learning algorithms when it exists. Some studies have been done in the literature, and methods have been proposed to deal with classroom noise. Zhu and Wu (2004) investigated the effect of classroom noise and variable noise separately and together on classification performance. They found that the existence of both problems separately and together negatively affected the classification performance. Brodley and Friedl (1999) proposed a noise identification approach for misclassified samples [8].

Another problem seen in general in health data sets is **outliers.** There are values well above or well below the optimal ranges in the data sets of the tests applied to patients to diagnose the disease. When the literature is reviewed, studies related to outlier observations have generally been related to the detection of outliers and their deletion from the data [9]. Deletion of data is undesirable, as it leads to loss of information. For this reason, it should be ensured that the contrary observations remain in the data set without being removed from the data. Bull et al. (2019) discovered outlier observations by using Mahalanobis distance in the method they proposed for outlier detection and feature extraction in high dimensional data [10].

In addition, health data sets contain an irrelevant variable that decreases or slows down the classification performance, as seen in other data [11]. Many feature selection methods have been proposed in the literature to remove these irrelevant variables from the data set. Yang et al. (1997) used genetic algorithms with 17 different data sets in feature selection methods in their study and showed that the classification performance increased in different algorithms [12]. In their study, John et al. (1997) solved over-learning by removing irrelevant variables from the data set with wrapper methods to overcome the over-learning problem in classification algorithms [13] Rodriguez et al. (2018) compared the classification performances of feature selection methods (filtering, wrapper, and embedded). They mentioned that filtering methods are faster than wrapper and embedded methods, which are slower but more successful than filtering methods [14].

This study aims to increase ensemble learning classification algorithms’ performance by removing the irrelevant variables from the data with the newly proposed feature selection method after eliminating the missing observation, class imbalance, and classroom noise problems that may arise in disease diagnosis data sets.

## 2. METHODS

The Pima, Parkinson, and SPEC Heart data sets used in the study were obtained from UCI and Keel machine learning databases [15]. While choosing data sets, it is essential to have different sample sizes, feature numbers, and at least one of the data preprocessing problems better to understand the effects of the algorithms in the study. In the study, considering the computer’s capacity as a calculation tool, data sets with low and different attribute numbers were preferred since larger sets cannot be studied. The data was divided into 70% training sets and 30% test sets when using classification algorithms. Among the ensemble learning algorithms, which are widely used in recent years and show high classification performance; Random Forest (RF) [16], Gradient Boosting Machine (GBM) [17], Extreme Gradient Boosting Machine (XGBM) [18], Light Gradient Boosting Machine (LGBM) [19], CatBoost [20], Bagging [21] algorithms were used. While calculating the classification performance of algorithms, among the measurement metrics; Accuracy (ACC), sensitivity (SEN), precision (PRE), F measure, kappa measure are used. In a statistical comparison of the T-score method and outliers’ accuracy values and correlation sensitive t-score (Outliers and Correlation t-Score, OCtS) methods’ accuracy values, the normality distribution assumption of the data was checked with the Shapiro-Wilk normality test. The Mann Whitney-U test was used to compare T-Score and OCtS methods. In the analysis, RStudio 1.2.1335, Python Jupyter Notebook 6.0.3, IBM SPSS Statistics 25 programs were used in the macOS-BigSur operating system. In all statistical analyzes, it was taken as α = 0.05.

### 2.1. DATA PREPROCESSING

#### 2.1.1 Missing Value

While doing research, missing values can occur for many reasons. Since each participant is essential for the study in research, loss of information is not desired. Therefore, with some statistical approaches, missing data can be completed in a meaningful way or deleted from the data set without distorting the significance. Missing value percentage can be calculated with the following formula (1) [22]:

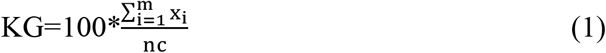

x_i_: number of independent variable missing values, n: number of observations, c:number of variables, i=1,…,m

#### 2.1.2 Class Imbalance

Class imbalance is the case where the total number of observations at the levels of the class variable differ from each other. The class imbalance problem occurs when a large class or majority class in a data set is significantly more than the other rare or minority classes. Undersampling methods make up a subset of the original data set as much as the minority class number by eliminating the observations of the majority class. Oversampling creates new examples from existing examples by replicating the minority class in the original data set as much as the majority class numbers. There are also hybrid methods combining both sampling methods. The most used method is the SMOTE algorithm, one of the resampling techniques. In this algorithm, each sample in the minority class is taken, and new samples are created synthetically by using the k-nearest neighbors algorithm [23]. The class imbalance rate (CI) formula (2) is as follows [24]:

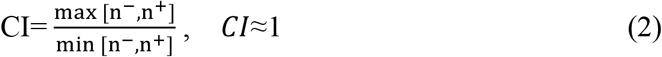

n^-^: one of the classes, n^+^: the other of the classes

#### 2.1.3 Class Noise

In most of the data used in research (except structured and synthetic environments), it is almost inevitable that the data contain noise. This “flawed data” can be caused by many factors such as faulty measuring devices, errors in data entry, transcription errors, or irregularities due to data transfer from different media. However, the accuracy of a classifier can be highly dependent on the quality of the data used during the training phase. For this reason, the results obtained from a noisy training set operated with the same classification algorithm may have less accuracy than those obtained from a noiseless data set [25]. The percentage of class noise (CN) is expected to be zero or close to zero and can be calculated by the formula (3) [26]:

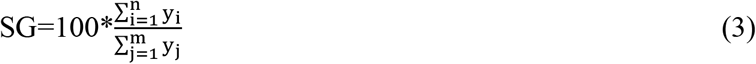

*y*_i_: total number of misclassified observations, *i*=1,…, *n*
*y*_j_: dependent variable total number of classes, *j*=1,…,*m*

### 2.2 FEATURE SELECTION METHODS

When the literature is reviewed, feature selection methods are grouped under three main headings [27]. The first of these is the filtering methods that are based on statistical methods and thus provide fast results. The second is wrapper methods that are based on machine learning methods and associate with the classifier at each stage. In the third group, there are embedded methods in which machine learning algorithms and feature selection methods work together and work by establishing a relationship with the classifier at each stage. While the feature selection is made, the following ranges can be used to decide the number of features required in selecting the threshold value of sorting methods according to the total number of features (N) [27]:

- If N≤10, choosing 75% of the features,
- If 10 <N <75, choosing 40% of the features,
- If 75 <N <100, selecting 10% of the features,
- If N≥100, it is considered appropriate to choose 3% of the features.

#### 2.2.1 Filter Methods

Filtering methods used in feature selection while reducing size is one of the oldest techniques [28]. Classification methods are not used in these operations using statistical methods. For this reason, algorithms work faster, and results are obtained faster. In this way, while making calculations, high benefits are provided in terms of time and cost. It has less processing complexity and higher explanation than the embedded and wrapper methods [27]. Commonly used filtering methods; chi-square test, Fisher score, t-score, Welch t statistics, knowledge gain, correlation-based filtering, Relief [29].

#### 2.2.2 Embedded Methods

In embedded methods, both feature selection algorithms and classification algorithms are used together. Therefore, embedded methods are slower and more costly than filtering methods like wrapper methods. Although filtering methods are fast and low cost, they do not use classification methods, so some problems or poor performance can be seen in classification. However, there can be an increased computational cost with the high dimensionality of coiled methods, particularly microarray data. Embedded methods have been discovered, which are an intermediate solution for researchers and use classification methods to generate criteria for sorting properties. For example, after creating and sorting subsets with decision trees or support vector machine methods, the desired level of features can be selected [27].

#### 2.2.3 Wrapper Methods

In Wrapper methods, the highest performance method is selected by using machine learning algorithms for feature selection. In this method, the best subset creation and selection techniques are more successful than filtering methods. However, considering the classifier’s success at each stage, it becomes slower and more costly. As subset search strategies, different methods such as consecutive forward selection, sequential backward selection, consecutive forward sliding selection, consecutive backward sliding selection, l add r exit, genetic algorithms can be used [27].

### 2.3 A NEW APPROACH

Statistical-based filtering methods are fast and low-cost, so their usability and explainability are better than other methods [30]. In the t-score method suggested in the literature, which is under the filter methods, outlier observations were not taken into account [31]. Today, contrary observations are found in many data, especially health data; Although it can be deleted from the data by an expert who created the data set or has information about it, this is not the desired situation as it will cause information loss. In addition, considering whether there is a correlation between the features in the data sets, it is expected that the attributes have a low correlation with each other but a high correlation with the class variable. For this reason, in the new approach, it is recommended not to delete outlier observations from the data set. The median approach, one of the robust techniques, was used in the t-score method, and the correlation approach suggested by Budak (2016) was added to the formula to show a low correlation between the features and high correlation with the class variable [29].

One of the methods used to detect outlier observations is the mean and standard deviation. In order to have descriptive information about data sets, mean and standard deviation values are used in continuous data as in the formula (4, 5) [32]:

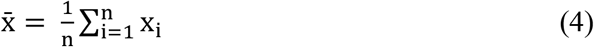

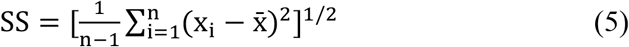

For example, when we look at Table 1 prepared from the data derived from the real values, the mean platelet in the blood of the patients is 177.50 while the standard deviation is 60.81. If the value 450 seen as an outlier is deleted, the average will change to 165.65 and the standard deviation to 18.56. Therefore, the presence of outlier observation negatively affects the mean and standard deviation. Under the assumption that the observation interval is [a, b], two standard deviations from the mean in the lower and upper range will be 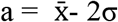 and 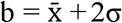, and observations outside these values will be defined as outlier. This situation can be used as ±1σ, ±2σ, and ±3σ. While the median of platelet values in the blood is 166.6, when the outlier observation value of 450 is deleted from the data, it becomes 165. As seen in the example, the median is not affected by the outlier observation as much as the average. The median approach can be used to avoid deleting outlier observations from the data [9]:

**Table 1.** Platelet counts in the blood

|  |  |  |  |  |  |  |  |
| --- | --- | --- | --- | --- | --- | --- | --- |
| 178 | 165 | 176 | 187 | 150 | 141 | 131 | 154 |
| 156 | 158 | 174 | 187 | 194 | 143 | 152 | 168 |
| 177 | 165 | 156 | 194 | 138 | 176 | 190 | 450 |

The data set is x_1_, x_2_, x_3_,…, x_n_; if n is odd, m=(n+1)/2

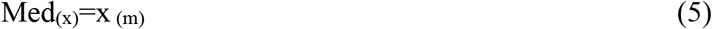

If n is even, m=n/2; The sample median is any point between x_(m)_ and x_(m+1)_, and is calculated by the following formula (6).

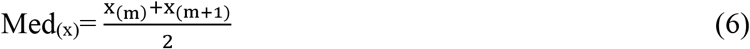

In data sets with outlier observations, the median is an excellent robust alternative to the mean, while the median of absolute deviations (MAD) is a good robust alternative to the standard deviation. MAD is defined by the following formula (7).

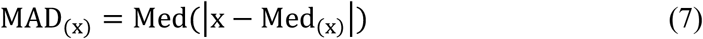

In order to compare the median of absolute deviations with the standard deviation, the normalized MAD, which is MADN defined by the formula (8) [9].

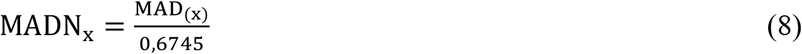

Based on this situation, using the median and MADN approach instead of the mean and standard deviation in the t-score method (9) for outlier observations is suggested. Thus, formula (10) was created for the feature selection method (Outliers t-Score, OtS) sensitive to outlier observations. The + and - signs in the formula represent different classes;

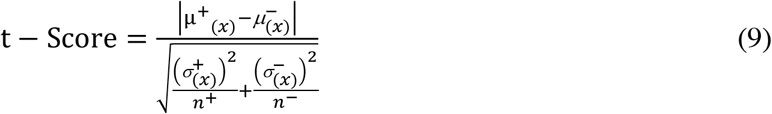

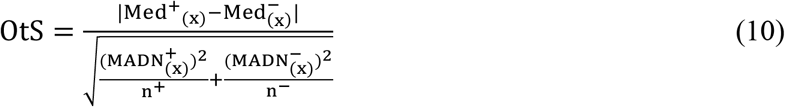

The correlation problem, which is generally seen in all data sets, still continues after the OtS method recommendation. Budak’s correlation approach is when |r_iy_|/r_iy_ is the correlation of the related attribute with the class variable riy and the mean of the internal correlation between the attributes 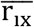 is added to the formula. The result of |r_iy_|/ r_iy_ will always be greater than 1, it will increase the chance of selection by raising the relevant feature higher in the ranking [29]. For this reason, the sensitive t-score (Outliers and Correlation t-Score, OCtS) method was recommended for outlier observations and correlation. In the correlation of the class variable (|r_iy_|), the absolute value is used in the formula (11) since the direction of the relationship is not essential. However, this method is recommended to be used when the number of classes is 2 and the median value of both classes is not zero.

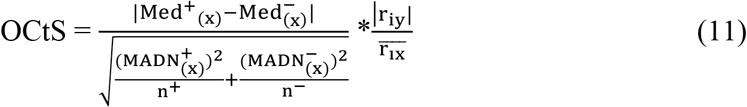

## 3. RESULTS

In this section, the problems that exist in the data sets are summarized, then the classification performance of the algorithms by solving these problems; The raw form of the data sets is compared with the data preprocessed form. Feature selection was made for OCtS and t-score method, and then the classification performances of the new data sets were compared. In addition, the statistical significance of this difference has been shown. Table 2 shows the problems of missing value (9.27%), class imbalance (1.86), and class noise (0.15%) in the Pima data set. Class noise (0.10%) and class imbalance (3.06) problems are observed in the Parkinson dataset, while there are no missing observations. There is only a class imbalance (1.61) problem in the SPEC Heart data set.

**Table 2:**
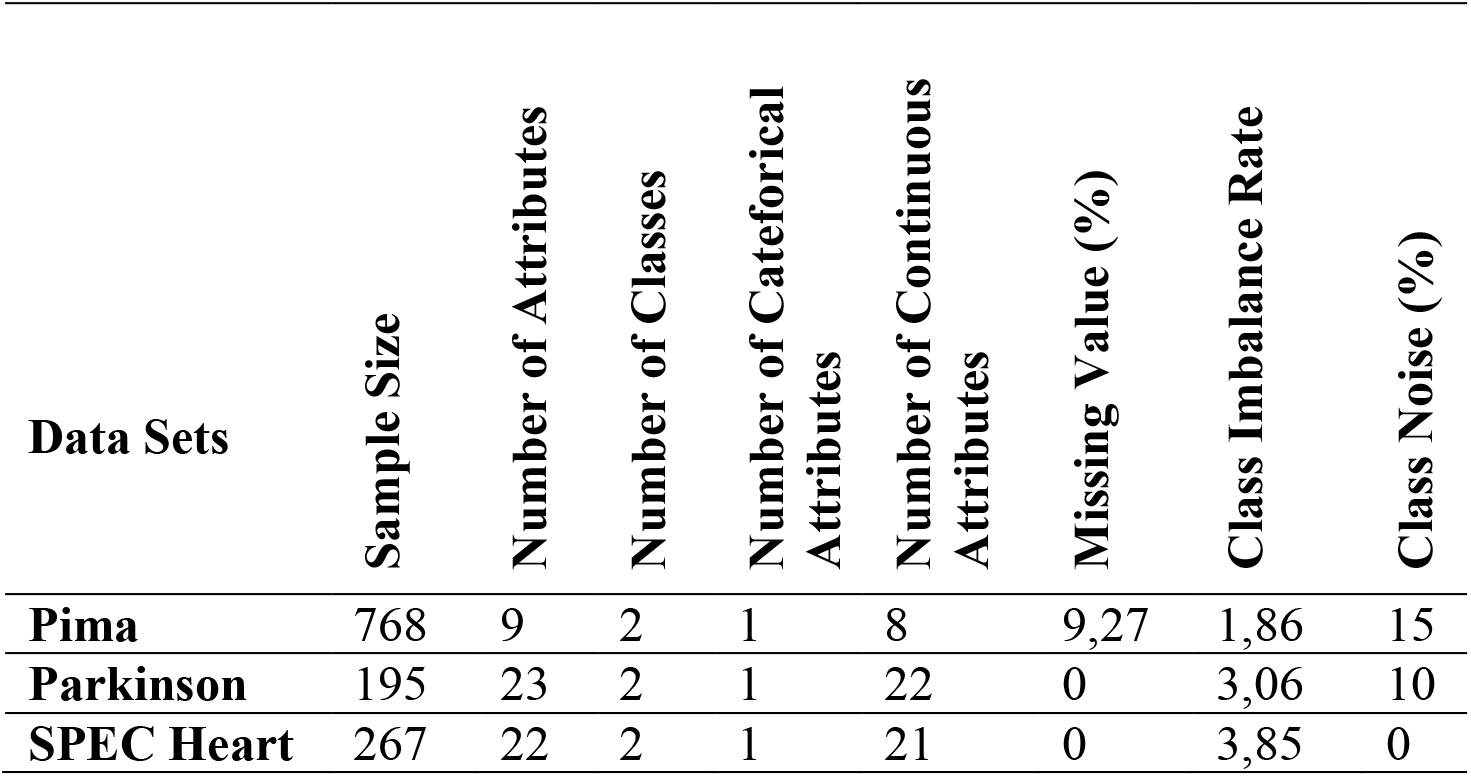
Information about data sets

Missing value assignment was performed under the assumption of MAR in the Pima data set with missing observations. Median and mean assignment methods were used to solve the missing observation problem. Considering the outliers in the variables, the median assignment was made for the continuous variables and categorical variables that were the outlier observations, and the mean assignments were made to the continuous variables that were not the outliers. The median assignment was made due to the outlier observation in Pres, Insulin, and Mass variables. The mean assignment was made to Plas and Skin continuous variables and since there were no outlier observations.

In the study, a class noise problem was observed in other data sets except the SPEC Heart data set. The hybridRepairFilter function from the NoiseFiltersR package was used in the R program to overcome the class noise problem. This function is preferred because it enables the relabeling of the observations without removing the observations with class noise from the data sets. The majority vote method was used for relabeling class noises. The last situation that occurred after the class noise problem in these data sets was solved was recorded as the new data set.

A class imbalance problem was observed in all data sets used in the study. This problem is solved with the SMOTE function in a program called Python. Here, by increasing the minority class as much as the difference between the number of classes, almost to the number of the majority class, the class numbers are equalized, and the class imbalance problem is solved. In Figure 1, class imbalance situations of data sets can be seen.

**Figure 1:**
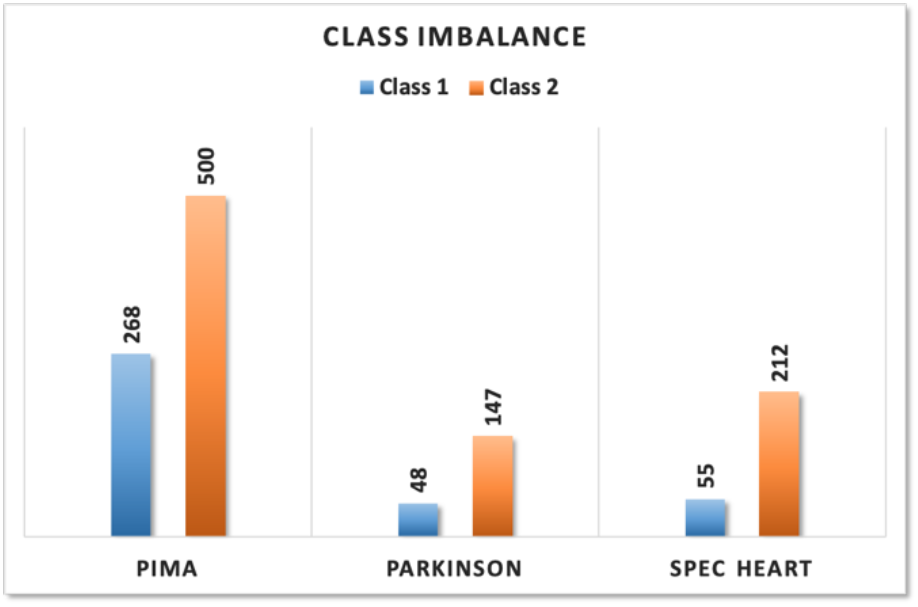
Class imbalance in datasets

In Pima data, the correlations of the features with the class are moderate for some variables and low for some variables. In the correlations between features, there is generally no high correlation are shown in Figure 2.

**Figure 2:**
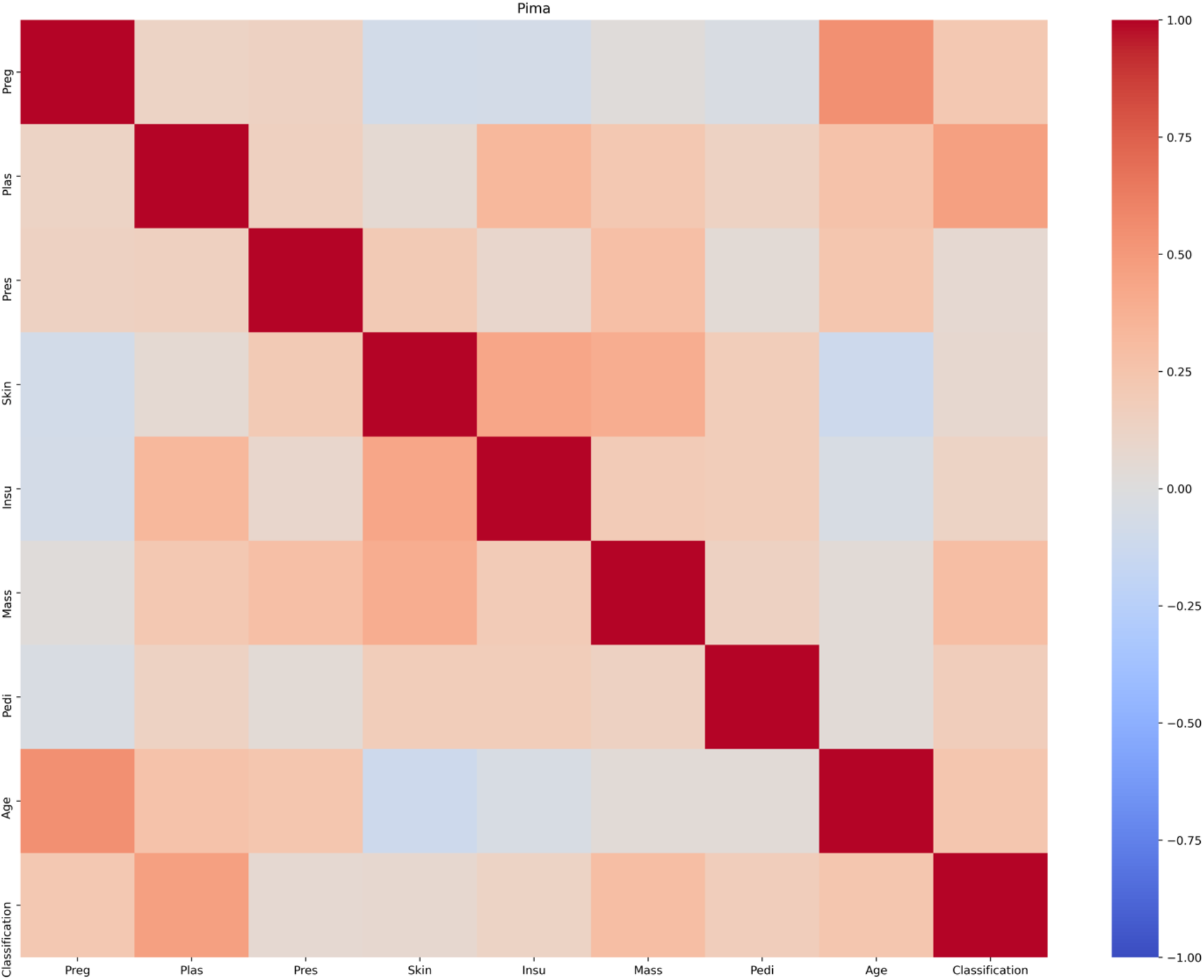
Pima dataset correlation heat graph

Looking at the temperature correlation graph for Parkinson’s data, the features show different degrees of positive and negative correlations with the class variable. There is a strong positive and negative correlation problem in some variables between the features are shown in Figure 3.

**Figure 3:**
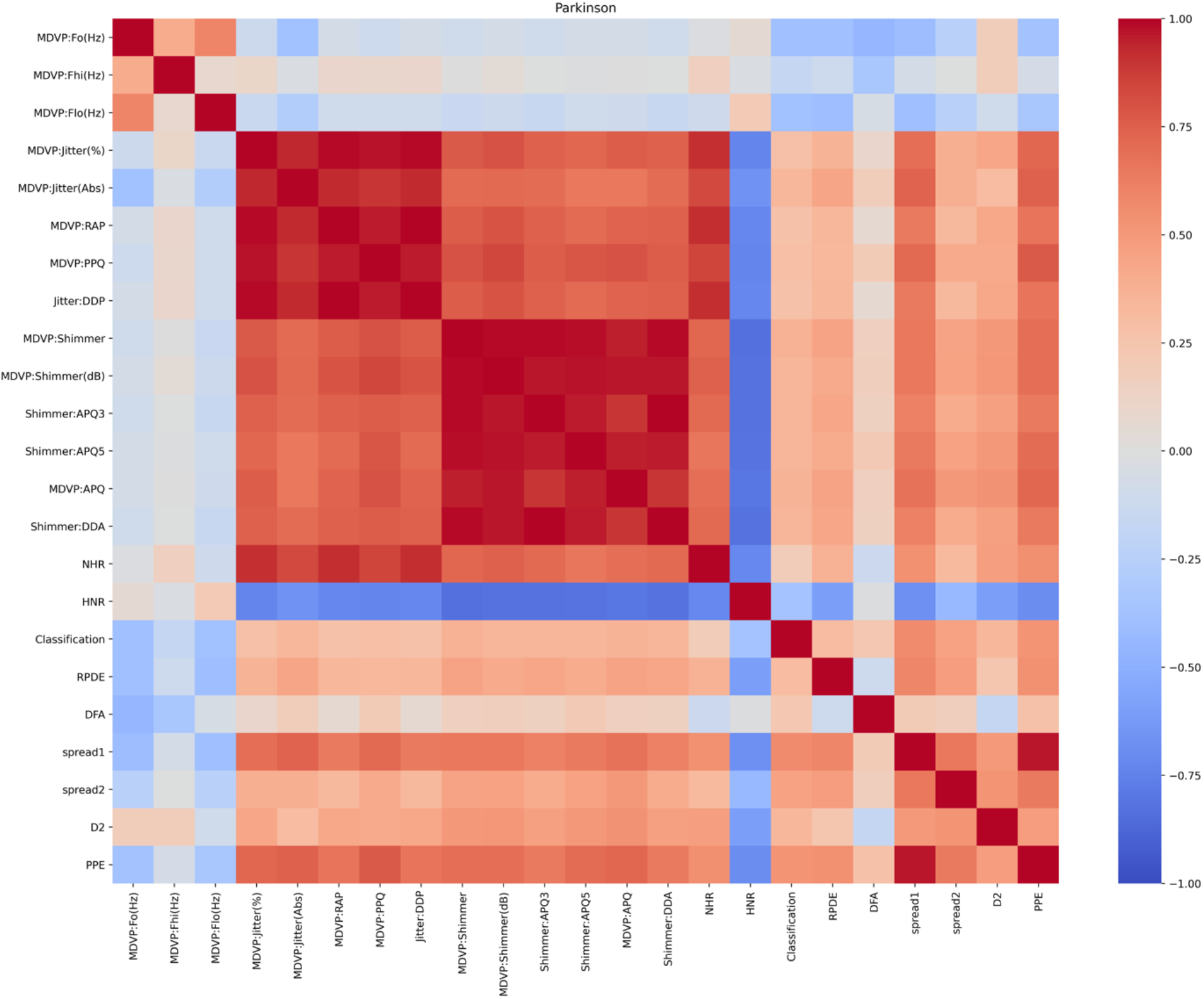
Parkinson dataset correlation heat graph

When the correlation temperature graph of SPEC Heart data is examined, it is seen that the correlations of the features with the class are generally negative in low or medium-strong. However, correlations between attributes are positive and strong for some variables are shown in Figure 4.

**Figure 4:**
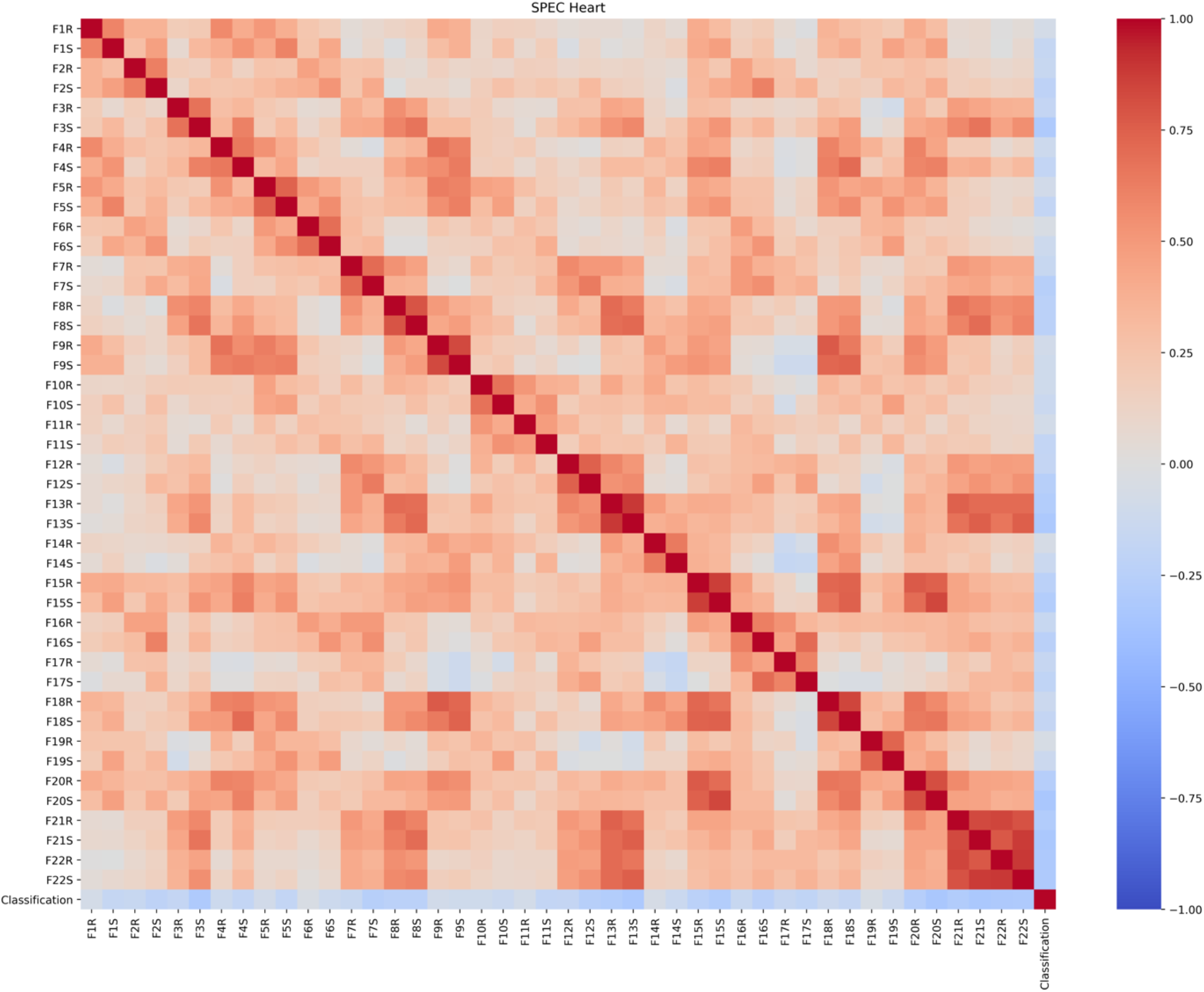
SPEC Heart dataset correlation heat graph

According to Figure 5, outlier observations of the first five attributes in the data sets are seen. Since the number of attributes in the data sets is much and creates confusion in the graphs, the first five variables were chosen randomly.

**Figure 5:**
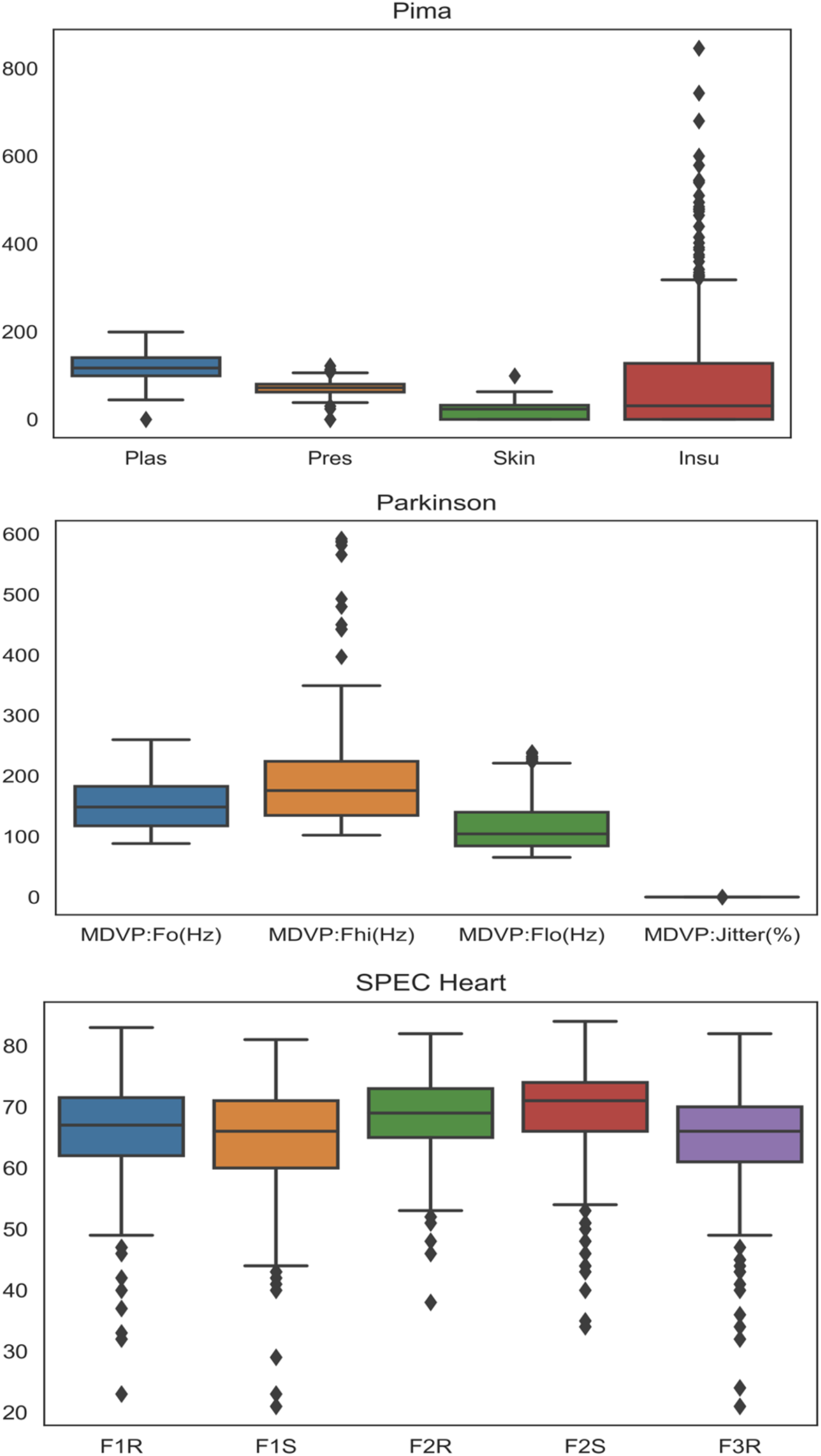
Top 5 variables and outlier observations in data sets

Table 3 shows the classification performances of the raw states of the data and the data preprocessed states. In all data sets and all algorithms, the data preprocessed state’s accuracy and kappa values were higher than the data’s raw state. The RF and Lightboost algorithms’ classification performance has reached an accuracy of 0.80 after the class imbalance, class noise, and missing value situation in the Pima data set have been eliminated. After correcting the class imbalance and class noise problems in the Parkinson dataset, it reached 0.91 percent accuracy in XGBM and Catboost algorithms. In the SPEC Heart data set, there is only a class imbalance problem. After the classes have been balanced, the classification performance has increased to 0.90 accuracies in RF and LightBoost algorithms.

**Table 3:** Classification performance of raw data and preprocessed data

|  |  | Raw Data |  |  |  |  | Preprocessed Data |  |  |  |  |
| --- | --- | --- | --- | --- | --- | --- | --- | --- | --- | --- | --- |
| Data Sets | Algorithm | ACC | SEN | PRE | F | Kappa | ACC | SEN | PRE | F | Kappa |
| Pima | RF | 0,65 | 0,66 | 0,67 | 0,65 | 0,24 | 0,80 | 0,81 | 0,81 | 0,81 | 0,61 |
|  | GBM | 0,66 | 0,66 | 0,68 | 0,68 | 0,27 | 0,78 | 0,78 | 0,79 | 0,78 | 0,56 |
|  | XGBM | 0,64 | 0,64 | 0,65 | 0,64 | 0,23 | 0,79 | 0,80 | 0,80 | 0,80 | 0,58 |
|  | LightBoost | 0,64 | 0,63 | 0,64 | 0,63 | 0,24 | 0,80 | 0,81 | 0,80 | 0,80 | 0,61 |
|  | CatBoost | 0,67 | 0,67 | 0,67 | 0,67 | 0,30 | 0,79 | 0,79 | 0,79 | 0,79 | 0,58 |
|  | Bagging | 0,62 | 0,63 | 0,63 | 0,63 | 0,15 | 0,76 | 0,77 | 0,77 | 0,76 | 0,52 |
| Parkinson | RF | 0,62 | 0,62 | 0,63 | 0,62 | 0,14 | 0,90 | 0,91 | 0,90 | 0,90 | 0,81 |
|  | GBM | 0,67 | 0,68 | 0,68 | 0,67 | 0,28 | 0,90 | 0,90 | 0,89 | 0,89 | 0,81 |
|  | XGBM | 0,67 | 0,67 | 0,68 | 0,67 | 0,29 | 0,91 | 0,91 | 0,91 | 0,91 | 0,90 |
|  | LightBoost | 0,69 | 0,71 | 0,70 | 0,70 | 0,31 | 0,89 | 0,90 | 0,89 | 0,89 | 0,79 |
|  | CatBoost | 0,66 | 0,67 | 0,67 | 0,66 | 0,22 | 0,91 | 0,91 | 0,91 | 0,91 | 0,88 |
|  | Bagging | 0,71 | 0,72 | 0,71 | 0,71 | 0,36 | 0,89 | 0,91 | 0,91 | 0,89 | 0,79 |
| SPEC Heart | RF | 0,80 | 0,82 | 0,82 | 0,82 | 0,30 | 0,90 | 0,90 | 0,90 | 0,89 | 0,82 |
|  | GBM | 0,79 | 0,80 | 0,79 | 0,79 | 0,31 | 0,89 | 0,90 | 0,90 | 0,89 | 0,81 |
|  | XGBM | 0,80 | 0,80 | 0,80 | 0,80 | 0,25 | 0,88 | 0,89 | 0,88 | 0,88 | 0,79 |
|  | LightBoost | 0,82 | 0,83 | 0,82 | 0,82 | 0,35 | 0,90 | 0,90 | 0,90 | 0,89 | 0,82 |
|  | CatBoost | 0,82 | 0,82 | 0,83 | 0,82 | 0,30 | 0,88 | 0,88 | 0,88 | 0,88 | 0,79 |
|  | Bagging | 0,78 | 0,79 | 0,79 | 0,78 | 0,32 | 0,84 | 0,86 | 0,85 | 0,84 | 0,70 |

In Figure 6, in the studies conducted on Pima, Parkinson, and SPEC Heart data sets, a new feature selection method sensitive to outliers and correlation (OCtS) showed higher accuracy in each algorithm and each feature number to the standard t-score method. In contrast, the RF algorithm shows the highest accuracy of 0.86 with six feature selection in Pima data. In the Parkinson data set, the GBM algorithm showed the highest accuracy of 0.98 with ten feature selection. SPEC heart data showed the highest accuracy of 0.98 with the XGBM algorithm with 14 feature selection.

**Figure 6:**
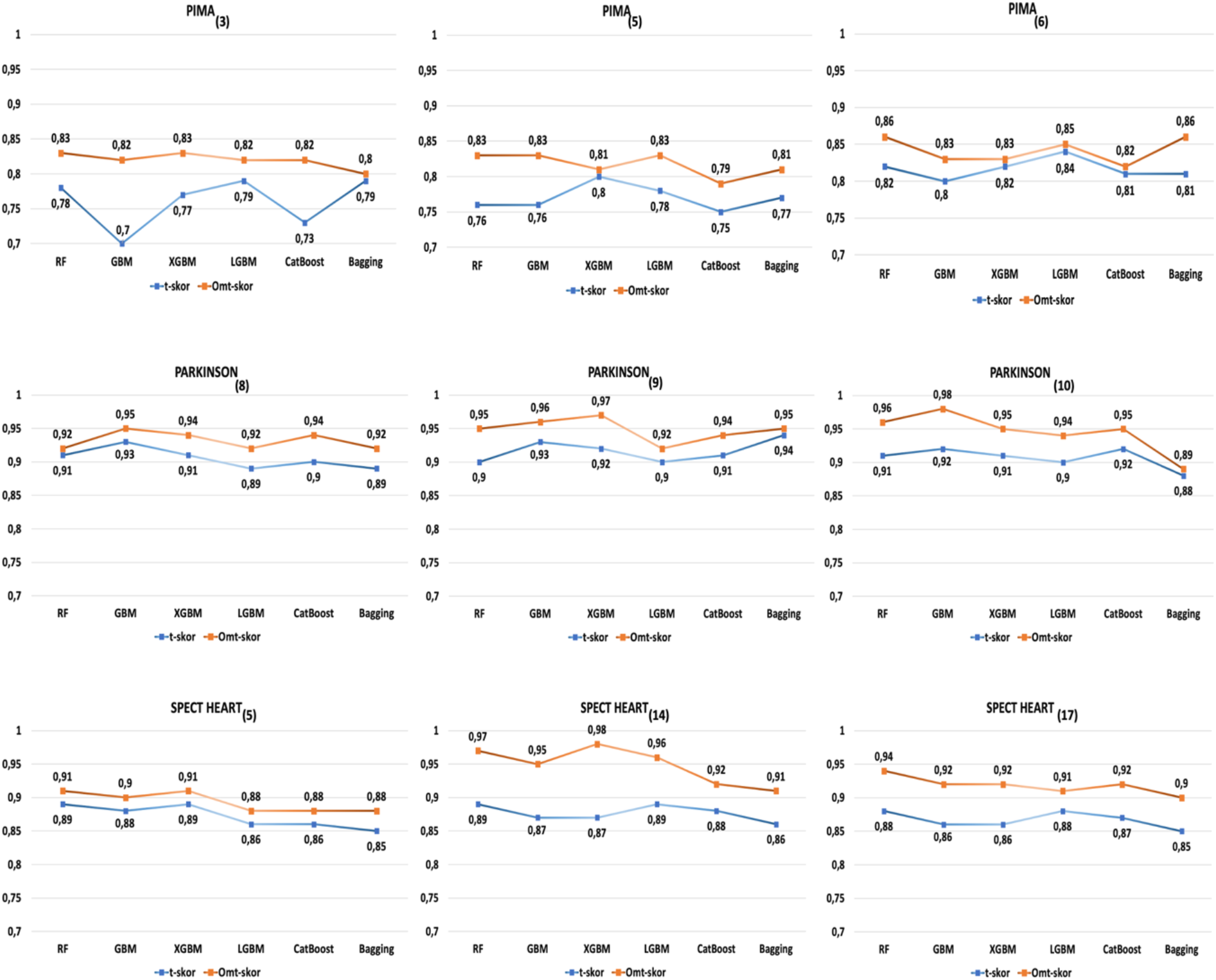
Accuracy percentages of t-score and OCtS methods after feature selection in all data sets

When looking at the findings related to the statistical comparison of the classification performances of the t-score method and the OCtS method, the accuracy values do not show normal distribution (p = 0.001 <α = 0.05). The difference between the classification accuracy percentages for t-Score and OCtS was statistically significant with 95% confidence (p = 0.001 <α = 0.05).

## 4. DISCUSSION

The algorithms’ classification performance in the raw state of the data sets used in the study showed very low values. This is due to the coexistence of some problems in data sets or one by one. These problems can be in the form of class imbalance, classroom noise, missing observation, outlier observation, and the correlation between independent variables. Besides, the fact that the number of variables and sample sizes are quite different from each other in the data sets negatively affects the classification performance. After eliminating these problems, the classification performance of the algorithms has increased considerably compared to the original classification performance. In the study, using the t-Score method, which is one of the feature selection methods used to solve the irrelevant variable problem in the data sets, a new feature selection method considering outlier observation and correlation situation is proposed. Classification performances of algorithms were compared for the T-Score method and OCtS method.

Considering the literature for Pima data, Akula et al. (2019) evaluated the classification performance of data preprocessing, feature selection, and ensemble learning algorithms in their study [33]. However, in this study, this problem was ignored by not mentioning any outlier observations existing in the data set, and the applications were made within the scope of outlier observation. Data normalization and missing data problems were focused on in the data preprocessing step class imbalance and class noise were not mentioned. Akyol et al. (2018) applied only data normalization among data preprocessing steps for Pima data; class imbalance, class appearance, and missing observation problems were not addressed, and in this study, the outliers in the data set were not mentioned [34]. They used Stability Selection (SS), Recursive Feature Elimination (RFE), and Iterative Relief (IR) methods for feature selection method. Researchers have preferred Adaboost, GBM, and RF methods from ensemble learning algorithms. According to the feature selection methods they use, the RF algorithm’s accuracy values are IR: 0.72, RFE: 0.71, SS: 0.73; The accuracy values of the GBM algorithm are IR: 0.70, RFE: 0.71, SS: 0.73. Our study showed that the RF algorithm showed a 0.86, and the GBM algorithm showed 0.83 accuracies of 0.83, outperforming these studies.

Looking at the literature for Parkinson’s data set, Patra et al. (2019) investigated the classification performance of simple and ensemble learning algorithms in their study [35]. The class imbalance problem in the data set was not corrected in the study, and no attribute selection was performed. When the ensemble learning algorithms’ classification performances are examined, 0.84 for Random Forest; An accuracy percentage of 0.81 was obtained for Bagging and 0.82 for Adaboost. Using Parkinson’s data set, Al Imran et al. (2018) investigated the effect of feature selection techniques on the performance of predicting Parkinson’s disease [36]. They solved the class imbalance problem with the SMOTE algorithm in the data preprocessing step. Then, as the feature selection method; They used Kruskal-Wallis Test, the Gain Ratio, Random Forest, Variable Importance, Symmetrical Uncertainty and RELIEF, methods and calculated the classification performance of the first 5, 10, 15, and 20 variables that they chose according to their importance. The first ten variables gave the highest accuracy percentage, 0.87, by the Random Forest algorithm and the Gain Ratio sorting method. In our practice, after all the problems were solved, and feature selection, the accuracy values of the algorithms were found to be higher than these studies in the literature; RF: 0.96; GBM: 0.98; XGBM: 0.97; LGBM: 0.96; CatBoost: 0.92; Bagging: 0.91.

When the literature is searched for the SPEC Heart data set, Madden (2002) presents an empirical comparison of The Markov Blanket Bayesian Classifier (MBBC) algorithm and three Bayesian classifiers in his study. In different data sets, Naive Bayes (NB) performances, Tree-Augmented Naive Bayes (TAN), and K2 algorithms were measured using accuracy and ROC curves. The class imbalance problem in the data set was not corrected in the study, and no attribute selection was performed. Therefore, the classification performance accuracy rate of algorithms; NB: 0.71; TAN: 0.81; K2: 0.80; MMBC: is in the form of 0.80 [37]. After the class imbalance was eliminated in our study and the feature selection was made with the proposed OCtS method, the XBGM algorithm showed the highest accuracy rate as 0.98, and a higher value was obtained from the existing studies.

In general, when looking at other studies in the literature, there is no study that evaluates all problems such as class imbalance, classroom noise, missing observation, irrelevant variable, outlier observation, and correlation, and makes a similar feature selection method our study.

## 5. CONCLUSION

For each data set used in the study, for each number of variables and each algorithm, the OCtS method showed higher performance than the t-Score method. Based on this, it can be said that the classification performance of the data that has been selected for attributes and does not have irrelevant variables is higher. Before using ensemble learning algorithms while classifying, data preprocessing steps should be applied, and then the appropriate attribute selection method should be applied to the data. In future studies, the proposed OCtS formula can be tested in larger data sets. The OCtS approach, which works only on 2-class data, can be expanded to be used for data sets with more than two classes.

## Notes

### Competing Interest Statement

The authors have declared no competing interest.

## REFERENCES

1. Rahul D. Machine learning in medicine. Circulation. 2015;132:1920–1930.

2. Lin J-H, Haug PJ. Data preparation framework for preprocessing clinical data in data mining. AMIA Annual Symposium proceedings AMIA Symposium. 2006:489–93.

3. Donders ART, van der Heijden GJMG, Stijnen T, et al. Review: A gentle introduction to imputation of missing values. Journal of Clinical Epidemiology. 2006.

4. Liu Y, Gopalakrishnan V. An Overview and Evaluation of Recent Machine Learning Imputation Methods Using Cardiac Imaging Data. Data. 2017;2(1):8–8.

5. Huang M-W, Lin W-C, Tsai C-F. Outlier Removal in Model-Based Missing Value Imputation for Medical Datasets. Journal of Healthcare Engineering. 2018;2018:1–9.

6. Leo J, Luhanga E, Michael K. Machine Learning Model for Imbalanced Cholera Dataset in Tanzania. Scientific World Journal. 2019;2019:1–12.

7. Seo JH, Kim YH. Machine-learning approach to optimize smote ratio in class imbalance dataset for intrusion detection. Computational Intelligence and Neuroscience. 2018;2018.

8. Brodley CE, Friedl MA. Identifying mislabeled training data. Journal of artificial intelligence research. 1999;11:131–167.

9. Maronna RA, Martin RD, Yohai VJ. Robust Statistics. Wiley; 2006. (Wiley Series in Probability and Statistics).

10. Bull LA, Worden K, Fuentes R, et al. Outlier ensembles: A robust method for damage detection and unsupervised feature extraction from high-dimensional data. Journal of Sound and Vibration. 2019;453:126–150.

11. John GH, Kohavi R, Pfleger K. Irrelevant features and the subset selection problem. Machine Learning Proceedings 1994: Elsevier; 1994. p. 121–129.

12. Yang J, Honavar V. Feature Subset Selection Using a Genetic Algorithm. Vol. 5. Boston, MA: Springer US; 1998. p. 117–136.

13. Kohavi R, John GH. Wrappers for feature subset selection. Artificial Intelligence. 1997;97(1-2):273–324.

14. Rodriguez-Galiano VF, Luque-Espinar JA, Chica-Olmo M, et al. Feature selection approaches for predictive modelling of groundwater nitrate pollution: An evaluation of filters, embedded and wrapper methods. Science of The Total Environment. 2018;624:661–672.

15. Machine Learning Repository. https://archive.ics.uci.edu

16. Breiman L. Random forests. Machine Learning. 2001.

17. Friedman JH. Greedy function approximation: a gradient boosting machine. Annals of statistics. 2001:1189–1232.

18. Chen T, Guestrin C, editors. XGBoost2016/08//; New York, NY, USA: ACM.

19. Ke G, Meng Q, Finley T, et al., editors. LightGBM: A highly efficient gradient boosting decision tree 2017.

20. Prokhorenkova L, Gusev G, Vorobev A, et al., editors. Catboost: Unbiased boosting with categorical features 2018.

21. Breiman L. Bagging predictors. Machine Learning. 1996;24(2):123–140.

22. Chen DY. Pandas for Everyone. 2017:161–161.

23. Chawla NV, Bowyer KW, Hall LO, et al. SMOTE: Synthetic minority over-sampling technique. Journal of Artificial Intelligence Research. 2002.

24. Vanhoeyveld J, Martens D. Imbalanced classification in sparse and large behaviour datasets. Data Mining and Knowledge Discovery. 2018;32(1):25–82.

25. Teng CM. Combining Noise Correction with Feature Selection. 2003. p. 340–349.

26. García S, Luengo J, Herrera F. Data Preprocessing in Data Mining. Vol. 72. Cham: Springer International Publishing; 2015. (Intelligent Systems Reference Library).

27. Bolón-Canedo V, Sánchez-Maroño N, Alonso-Betanzos A. Feature Selection for High-Dimensional Data. Cham: Springer International Publishing; 2015. (Artificial Intelligence: Foundations, Theory, and Algorithms).

28. Deng X, Li Y, Weng J, et al. Feature selection for text classification: A review. Multimedia Tools and Applications. 2019;78(3):3797–3816.

29. Budak H, Erpolat Taşabat S. A MODIFIED T-SCORE FOR FEATURE SELECTION. ANADOLU UNIVERSITY JOURNAL OF SCIENCE AND TECHNOLOGY A - Applied Sciences and Engineering. 2016;17(5):845–845.

30. Zhao XS, Bao LL, Ning Q, et al. An Improved Binary Differential Evolution Algorithm for Feature Selection in Molecular Signatures. Molecular Informatics. 2018;37(4):1700081–1700081.

31. Wang S, Li D, Song X, et al. A feature selection method based on improved fisher’s discriminant ratio for text sentiment classification. Expert Systems with Applications. 2011.

32. Hastie T, Tibshirani R, Friedman J. Elements of Statistical Learning 2nd ed. 2009.

33. Akula R, Nguyen N, Garibay I, editors. Supervised Machine Learning based Ensemble Model for Accurate Prediction of Type 2 Diabetes2019/04//: IEEE.

34. Akyol K, Şen B. Diabetes Mellitus Data Classification by Cascading of Feature Selection Methods and Ensemble Learning Algorithms. International Journal of Modern Education and Computer Science. 2018;10(6):10–16.

35. Patra AK, Ray R, Abdullah AA, et al. Prediction of Parkinson’s disease using Ensemble Machine Learning classification from acoustic analysis. Journal of Physics: Conference Series. 2019;1372:012041–012041.

36. Al Imran A, Rahman A, Kabir H, et al. The Impact of Feature Selection Techniques on the Performance of Predicting Parkinson’s Disease. International Journal of Information Technology and Computer Science. 2018;10(11):14–29.

37. Madden M. Evaluation of the Performance of the Markov Blanket Bayesian Classifier Algorithm. arxivorg. 2002.

